# Intricacies of running a route without success in night-active bull ants (*Myrmecia midas*)

**DOI:** 10.1101/2024.12.10.627864

**Authors:** Sudhakar Deeti, Muzahid Islam, Cody Freas, Trevor Murray, Ken Cheng

## Abstract

How do ants resolve conflicts between different sets of navigational cues during navigation? When two cue sets point to diametrically opposite directions, theories predict that animals should pick one set of cues or the other. Here we tested how nocturnal bull ants *Myrmecia midas* adjust their paths along established routes if route following does not lead to their entry into their nest. During testing, foragers were repeatedly placed back along their homeward route up to nine times, a procedure called rewinding. This procedure produced an accumulating path integrator, or vector, in diametric opposition to the learned landmark views of the route. Repeated rewinding made some individuals head initially in the nest-to-feeder vector direction, but all ants ended up using the visual scene for homing, demonstrating the importance of view-based homing in this species. Repeated rewinding, however, led to path deteriorations, with increased path meander and scanning, results also found in desert ants. After nine rewinding trips, ants were displaced off their route in further manipulations, to a site near the nest, an unfamiliar site, or with the terrestrial surround entirely covered. The results show that a change in visual conditions diminished the weight accorded to path integration: the off-route ants no longer headed off in the vector direction as they did on the immediately preceding trial. They relied on celestial compass cues in other ways for homing. Experiment 2 showed the effects of rewinding in the unaltered natural habitat were not view-specific in these bull ants.

## Introduction

The remarkable abilities of visually navigating insects illustrates that even animals with small brains and low-resolution visual systems can still produce efficient and robust navigation in complex environments (Baddeley, Graham, Husbands, & Philippides, 2012; Warrant, 2017; Freas and Cheng 2022). As these animals explore their world, they learn to traverse specific paths leading to important locations, such as food sites or their nest. For example, nocturnal bull ant *Myrmecia midas* foragers follow stereotypical routes to and from a specific foraging tree and their nest. Such paths are idiosyncratic, indicating that ants are driven by personally acquired cues rather than by chemical pheromones or by interaction with other foragers (Freas, Narendra, & Cheng, 2017; Freas, Narendra, Lemesle, & Cheng, 2017; Islam, Freas, & Cheng, 2020). Indeed, solitary ant foragers show the ability to reach desired locations using multiple navigational systems, multiple sources of information, and through a variety of fine-scale behaviors. In particular, these ants rely on three main forms of navigational strategies, path integration (PI), view-based navigation, and a backup strategy called systematic search (Ferdous et al. 2018).

The well-studied phenomenon of PI is a strategy through which animals continuously combine odometric and compass information into a vector, a directional and distance estimate of their start location (Wehner & Srinivasan, 2003). When traveling, the animals combine their compass information with estimates of speed or distance constantly to update a home vector, which they can use at any time to guide them directly back to their starting point of journey, typically the nest (Collett, Collett, Bisch, & Wehner, 1998; Haferlach, Wessnitzer, Mangan, & Webb, 2007; Webb, 2019). Ants living in landmark-rich environments also learn and use the information provided by the visual panorama to facilitate navigation, whether they are diurnal (Cheng, Narendra, Sommer, & Wehner, 2009; Deeti, Fujii, & Cheng, 2020; Wehner, Michel, & Antonsen, 1996; Wystrach, Beugnon, & Cheng, 2011, 2012) or nocturnal foragers (Freas et al., 2017; Islam et al., 2020; Reid, Narendra, Taylor, & Zeil, 2013; Warrant & Dacke, 2011). When ants are uncertain about the nest position, or in the absence of PI and learned visual cues, foragers employ systematic search as a back-up mechanism, in which they perform looping search walks until they find information, whether local, panoramic, or celestial, which will aid navigation ( Ferdous et al. 2018; Schultheiss & Cheng, 2011; Schultheiss, Wystrach, Legge, & Cheng, 2013; Schultheiss, Cheng, & Reynolds, 2015; Wehner & Srinivasan, 1981).

Ants combine the multiple sources of navigational information flexibly. During the first inbound journey, ants use PI-based guidance and goal-directed image-matching information acquired near the goal (Collett, 2012). The route memories then produce precise paths even in the absence of a celestial compass (Collett, 2012; Collett, Collett, Chameron, & Wehner, 2003). Experiments that put two sources of information in conflict show that ants in general combine the two guidance systems in steering a path (Collett, 2012; Bregy, Sommer, & Wehner, 2008). In some circumstances, foragers appear to choose the optimal Bayesian weighting of different information sources, with the weighting assigned to a source dependent on its reliability (Legge, Wystrach, Spetch, & Cheng, 2014; Hoinville &Wehner, 2018; Narendra, 2007; Wehner, Hoinville, Cruse, & Cheng, 2016; Wystrach, Mangan, & Webb, 2015). Thus, all systems operate simultaneously, and their outputs are integrated, depending on the certainty and reliability of sources and on the motivational state of the animal.

Other flexibly adjusted behaviors characterize the ants’ navigational trips. Ants oscillate from side to side when they walk home (Cheng, 2022; Clement, Schwarz, & Wystrach, 2022; LeMoel & Wystrach, 2020; Murray et al., 2020; Wystrach, Buehlmann, Schwarz, Cheng, & Graham, 2020). In uncertain conditions or facing conflicting cues, foragers paths become less straight with individuals zig-zagging more from side to side during homing, a behavior called meandering (Wystrach, Schwarz, Schultheiss, Beugnon, & Cheng, 2011; Wystrach et al., 2014). In uncertain conditions, ants also stop forward movement and scan the environment more frequently. When views become unfamiliar because experimenters had manipulated the natural scene (Wystrach et al., 2014) or when learned views become aversive because experimenters set a pit trap on the route (Wystrach et al., 2020; see also Le Moël & Wystrach, 2020), ants meander more and scan the panorama more often (Freas, Wystrach, Schwarz, & Spetch, 2022; Zeil, Narendra, & Stürzl, 2014).

One way to introduce a conflict between different navigational systems and inject uncertainty is an experimental manipulation called rewinding. A homing ant nearing its nest is captured and placed back somewhere on its homebound route (e.g., the starting point, such as a feeder) and allowed to home again, re-traversing its previous route (Wystrach, Schwarz, Graham & Cheng, 2019). With repeated rewinding, the rewound ant’s path integrator continues to accumulate, creating an nest site estimate in the opposite, nest-to-feeder (outbound) direction. Each additional run the forager completes increases this conflict by extending the PI estimate distance in the opposite direction. Being captured and placed back on the route again also means a lack of success in homing, meaning there is a lack of reinforcement learning of the inbound route being associated with successfully reaching the nest and at least a form of extinction and perhaps also an aversive event tied to views leading up to this collection. All these factors should work to reduce the confidence and willingness to use the view-based cues that, as always, point from anywhere on the route to the ant’s nest. Given these changes in reinforcement and view memory valence, we would expect behaviors characteristic of being in uncertain situations should increase after rewinding. Empirical evidence so far supports these predictions.

In *C. bicolor*, ants were rewound repeatedly in a narrow channel surrounded by visual landmarks. Subsequently, in the absence of familiar visual landmarks around the channel, they dashed off in the opposite, nest-to-feeder direction (Andel & Wehner, 2004). Without the usual route-specifying landmark cues, the ants followed their vector, which, after repeated rewinding, pointed in the opposite direction from home. *Cataglyphis fortis* ants were trained along a route that passed a single dominating landmark in a visually barren salt-pan environment and were then rewound once. With the landmark still in place, they displayed what was called “a period of apparent confusion” on their next trip (Collett, 2014, p. 1), suggesting that a single unsuccessful event reduces the ants’ willingness to rely on their visually defined route. Independently of the number of times they had been picked up, *M. bagoti* foragers that had accumulated a longer conflicting PI vector in the nest-to-feeder direction showed more meandering and pausing to scan (Wystrach et al., 2019). With repeated rewinding, ants eventually made U-turns, shifting to the opposite direction.

Wystrach et al.’s (2019) study also showed that ants were particularly affected by rewinding during the portion of the journey in which they were rewound, suggesting view specificity in the effects of rewinding. Thus, rewinding has the most effect in the particular context in which the rewinding event took place, a context effect on learning that is found in other situations in insects (honeybees: Cheng, 2005; bumblebees: Colborn, Ahmad-Annuar, Fauria, & Collett, 1999) and vertebrate animals (Bouton, Rosengard, Achenbach, Peck, & Brooks, 1993; Bouton, Woods, & Pineño, 2004).

Context effects are also found in the path integration system. Desert ants run a shorter distance of their PI vector when the test environment in which they are running is dissimilar to the environment in which the vector was accumulated (Cheng, Middleton, & Wehner 2012; Cheng, Schultheiss, Schwarz, Wystrach, & Wehner, 2014; Narendra, 2007). We tested context specificity of the rewinding manipulation in night-active bull ants as well as the effects of rewinding when ants are displaced to different locations off their usual route after a number of rewound trips.

The test animals, *Myrmecia midas*, live in a visually rich environment and are known to rely heavily on terrestrial visual cues, though also attend to the visual cues of the overhead celestial compass, a key component of path integration (Freas, Narendra, & Cheng, 2017). Here, we rewound these nocturnal bull ants 9 times so that they completed the homeward route 10 times without entering the nest. We examined whether these rewound foragers showed any changes in their path straightness, scanning, and U-turns (a term we use to define foragers initially orienting in the direction opposite to the nest for a short distance before reorienting in the nest direction). With repeated re-runs, we predicted an increase in path degradation, lower path straighness with more meandering, scanning, and U-turns. After two rewinds in this experimental setup, the vector from path integration begins to point in the direction opposite of the true nest. After these rewound trips, ants were tested once after being displaced to locations off their route home. Ants were tested with a view from a location near the nest but off their usual route, a view off their habitual route some distance from the nest, and no useful view at all. With all these displacements following rewinding, we expected less reliance on path integration in the ants’ initial headings because the context had changed, especially in this species that is so reliant on using views. A view change means a context change from the context in which the path-integrated vector was acquired, making the ant less willing to use path integration (Cheng et al., 2012). It is difficult, however, to predict what actual direction they would head in. As with *M. bagoti* (Wystrach et al., 2019), we predicted context specificity in the effects of rewinding. The effects of rewinding should be larger in the segment in which the ants were rewound.

## General Methods

### Animals

The experiments were conducted on one nest on the northern portion of Macquarie University Wallumattagal campus in Sydney, Australia (33°46’11’’ S, 151°06’40’’ E), from November 2019 to March 2020 and control conditions were conducted from January 2021 to February 2021. The landscape around *M. midas* nests at our test site consisted of grass, *Eucalyptus* trees, litter, and leaves on the open ground. The ants usually nest close to a *Eucalyptus* tree (< 30 cm). Some portion of the ants forage on the tree located at the nest, called the nest tree, during the evening twilight. Workers also forage on trees around the nest; most of the ants forage at a nearby tree, called a foraging tree (Freas et al., 2017). They perform a single foraging trip for the night on the preferred tree, stay the night in the tree canopy before returning to the nest in the pre-dawn twilight. We observed that foragers made four to five foraging forays per week in the southern summer period (December to February). Each forager traveled to the same foraging tree each night. *M. midas*’ foraging activity started just before sunset, and in the evening twilight, ants came out of their nest.

From the chosen nest, the ants that were foraging on a particular tree 8 m away were captured in the evening twilight between 8:00 p.m. – 9:00 p.m. at the base of their foraging tree trunk, using foam-stoppered transparent plastic tubes. Ants were provided with a small amount of honey water and kept overnight in the laboratory. For adequate visibility during testing, all the experiments were conducted after sunrise on the following day, 6:30 a.m. – 10:30 a.m.

We tested each individual ant only once. To avoid the repeated testing of individual foragers, ants were painted after testing. Ants were kept it in an ice box for 10 mins to cool them down and then marked with yellow color using Citadel Paint™. After painting, ants were transferred back into the tubes. Within 2 to 3 mins they resumed normal activity and were released at the nest entrance. While Australia does not have ethical requirements concerning working with ants, we used experimental procedures that were minimally invasive and caused no notable adverse effects on the individuals or their colony (Freas et al., 2017).

### Experimental setup

The foraging route corridor between the nest and foraging tree contained tree mulch on the ground with sparsely distributed vegetation. Before the experimentation began, we cleared the sparsely distributed vegetation on the ants’ foraging corridor to ensure visibility while following homebound foragers. Then, we measured the distance between the nest and the foraging tree (8 m). We habituated ants to a white colored wooden plank (2 m × 1.5 m × 2.5 cm) by placing it at ground level near the foraging tree a month before the start of experimentation and allowed foraging ants to freely pass over this board each night during foraging. This plank provided a high-contrast background to obtain precise tracks of the foragers during experiments. We made a 50cm radius goniometer, with a circular indentation in the center with a 15mm diameter and 25mm depth to release the ants into (Deeti & Cheng, 2021; Deeti, Fujii, & Cheng, 2020). This central indentation on the surface of the recording area allowed us to relase foragers into this recess, preventing them from immediately bolting in a random direction. The distance from the center of the platform to the foraging tree was 1 m, thus 7 m from the nest. The distance from the wooden plank to the nest was further divided into two divisions called Section-A and Section-B, each 3.5 m (Figure 1). In Section-A, we constructed two parallel barriers of silver metal 3.5 m in length and 8 cm in height placed adjacent to the 1m white string grid on the ground (Islam, Deeti, Kamhi, & Cheng, 2021a). The foragers were habituated to the metal bars coated with ducting tape for several weeks along with the wooden board. The barriers helped to demarcate the end of section-A as well as to guide ants to travel along the foraging corridor.

**Figure 1.**
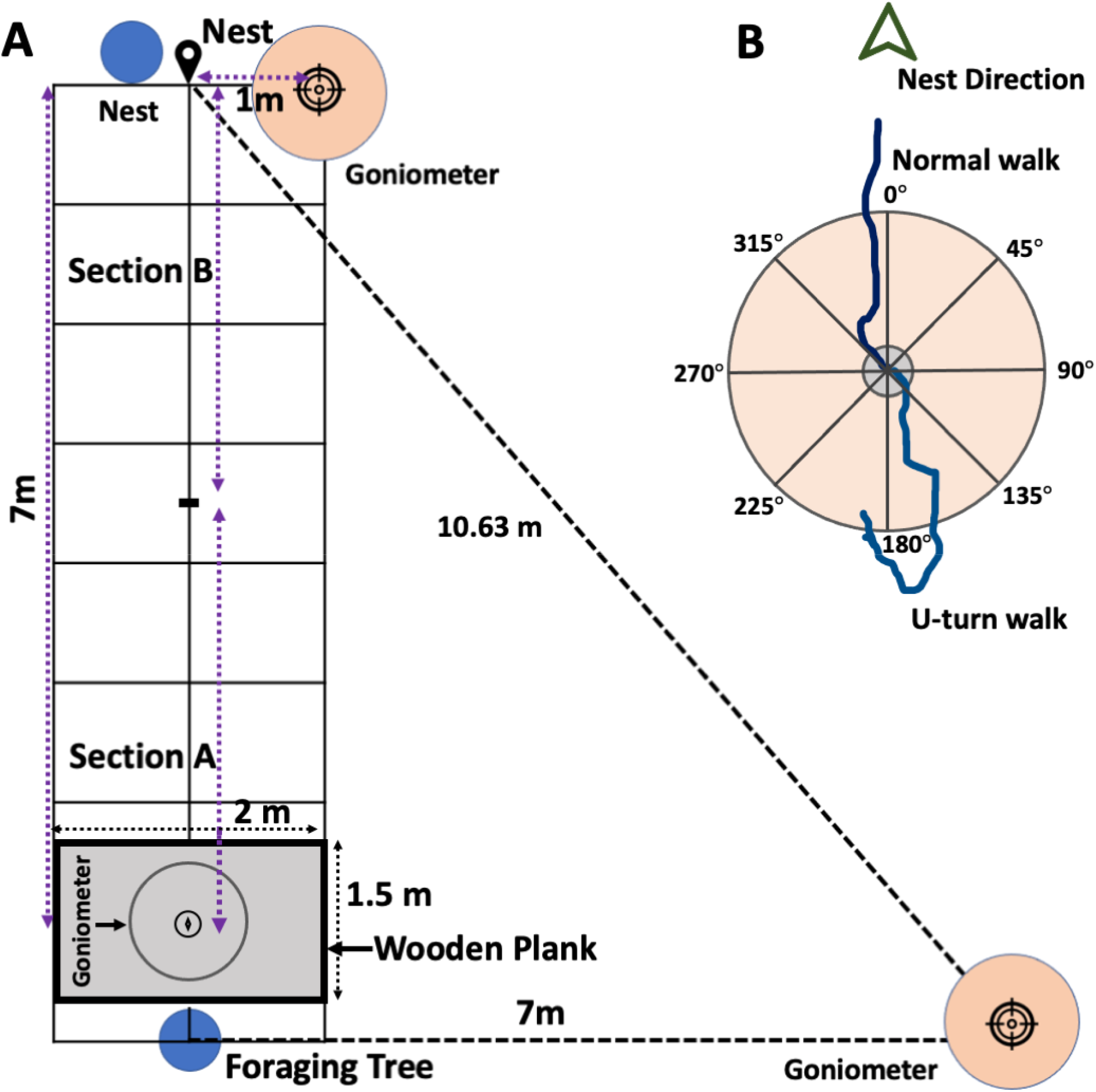
The experimental set up for the rewinding experiments. (A) The ants’ homing corridor was gridded with string between the nest and foraging tree. The blue circles indicate the tree positions and the larger circles with pointers indicate the release points for the Nest panorama (topmost circle), Unfamiliar panorama (rightmost circle), and Covered panorama tests. (B) The exemplary paths of one ant’s normal and U-turn paths starting on the goniometer.

### Testing and data recording

One experimenter manipulated the ant while a second observer recorded the ant’s path on graph paper. The two experimenters (SD and MI) switched roles frequently in the course of the study. Ants were placed in the indentation at the center of the goniometer for testing. To quantify the compass direction of their initial headings, the foragers’ first crossings at 50 cm and the corresponding 15° goniometer wedge, were noted. In Experiment 1, any U-turns moving in the opposite, nest-to-foraging-tree direction were noted, whether at the start of the trip or somewhere along the way. (It turned out that all such U-turns were at the beginning of a trip.) An ant was considered to have made a U-turn if it headed off the goniometer in a 45° sector around the nest-to-foraging-tree direction (Ants did not U-turn at all in Experiment 2). The number and positions of scans were also recorded in both experiments. A scan consisted of a head rotation between 1° and 360° of angular orientation followed by a pause to look direction of at least 2 seconds.

To obtain interrater reliability for the paths and scans, the two experimenters independently recorded the path coordinates of 10 ants not in any experiment for a 1-m distance. The ants were kept overnight as in the experiments, and they were returning home the next morning in daylight. This exercise was carried out after the main experiments were done. Independent scoring means that the two recorders were observing and recording the same ants simultaneously, but not observing how the other observer was drawing. (Indeed, it would have been impossible to draw an ant’s path should one observer focus on another observer rather than the moving ant.) The location on the *x*-axis at each 10 cm on the *y*-axis (y = 10. 20, etc.) were recorded. The Intra-class correlation coefficient (ICC) indicated a fair degree of reliability between two observers on the path coordinates and on the number of scans (paths: *ICC* = 0. 94, (*k*) = 0.74, *p* < 0.0001; scans *ICC* = 0. 96, (*k*) = 0.56, *p* < 0.0001), indicating that the intra-class correlation coefficients and Kappas (*k*) were significantly different from zero. To determine if the means of path straightness and scan rate differed across rewinding trials, we used a one-way repeated-measures ANOVA. A quadratic parametric regression model used to fit the dependent measures of path straightness and scan rate as a function of rewinding trials.

### Experiment 1: Full-path rewinding

The aim of the Full-path rewinding experiment was to test the effects of repeated rewinding along a homeward route on the night-active *Myrmecia midas*. Ants were rewound 9 times during their homeward route and we recorded the initial headings of reword ants as well as path straightness. After 9 rewound trips, ants were tested once at on-route or off-route locations to examine the effects of rewinding. In the corresponding control treatments, ants were allowed once to reach the nest area and then displaced once to the same set of locations. All control treatments were carried out on naive ants belonging to a different cohort from the rewound ants.

## Methods

Outbound foragers were collected at the base of their foraging tree during their outbound journey, fed, and kept overnight, which ensured they had a strong motivation to return to the nest. In the morning, ants were released at the foraging tree and allowed to return to their nest, in so doing winding down their PI vector to near zero. Each rewind then means recollecting the ants as they approached the nest (< 30 cm) and displacing them back on the route near the foraging tree, each rewind causing their PI vector to accumulate in the opposite direction to the traveled direction, that is, in the nest-to-foraging-tree direction. In each rewind, an ant near the nest was collected in a test tube and immediately brought back to the center of the goniometer placed at the center of the displacement platform. Nine rewind cycles in a row (resulting in inbound 10 trips) were performed on each ant (N =75). U-turns and the number and positions of scans were noted. Initial headings of the ants on rewinding trials 1, 5, and 9 (trips 2, 6, and 10) were noted for later analysis.

### On- and Off-Route test conditions

After completing 9 rewinding trials, ants were subjected to one of the on-route and off-route manipulation conditions: Nest panorama test (NPT), Covered panorama test (CPT), Unfamiliar panorama test (UPT) conditions. Control ants from a separate cohort were tested in corresponding conditions without having been rewound multiple times. In the Nest panorama (NPT) condition, ants (N = 25) were displaced 1 m from the nest in a direction perpendicular to the foraging-tree-to-nest direction (Figures 1 and 2). In the Covered panorama condition (N = 25), a 1m × 1m area around the wooden releasing platform was blocked with 1.5-m-high black, opaque sheets on four sides. In the Unfamiliar panorama condition, 25 ants were tested off-route with the open panorama (Figures 1 and 2), displaced 7 m from the foraging tree in a direction perpendicular to the foraging-tree-to-nest direction. The goniometer was placed on the ground. Initial headings on the goniometer were noted, but we did not record the paths from the displacement sites to the nest. Instead, the test ants were collected and released at the nest site to return home.

**Figure 2.**
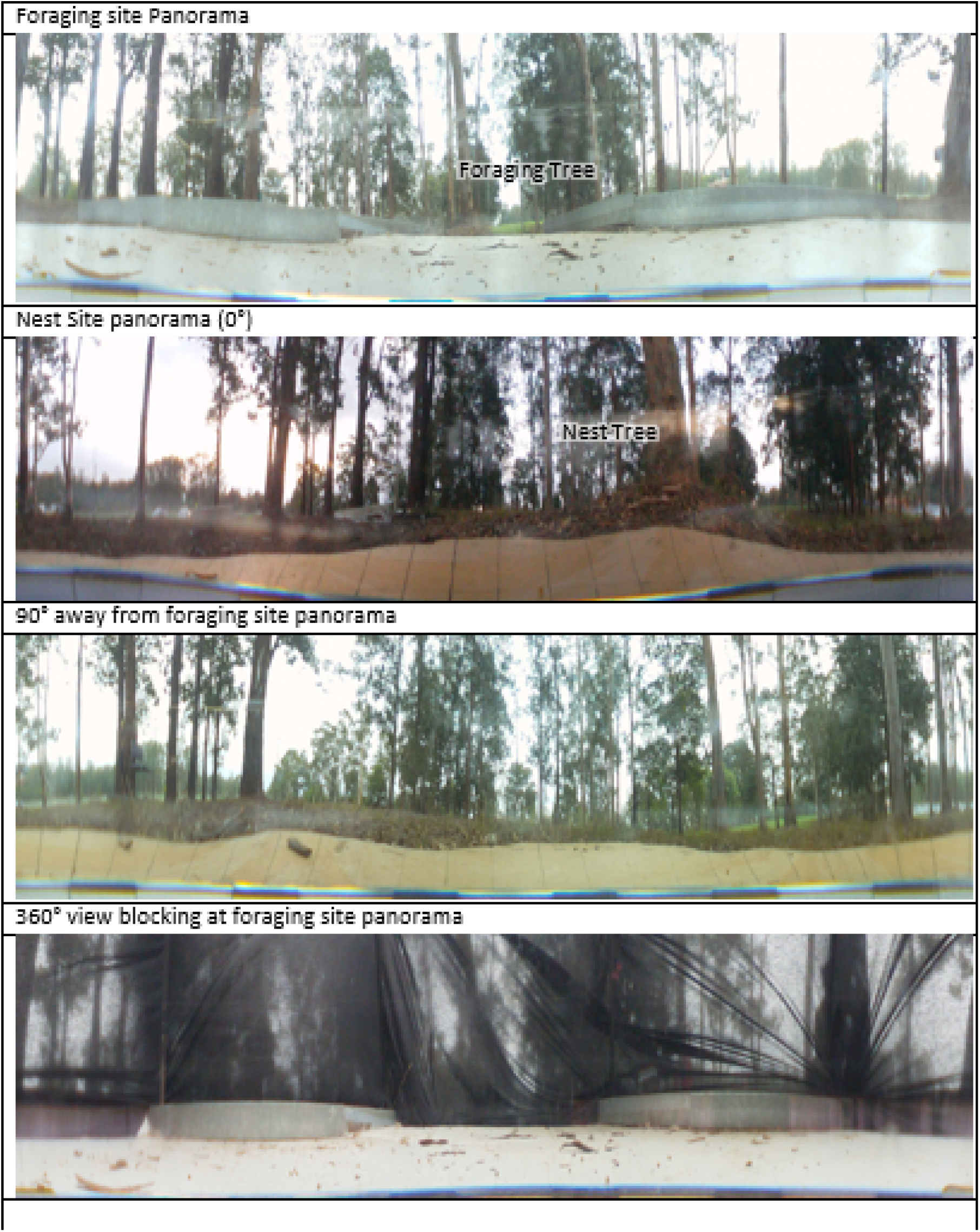
Panoramic 360° images of the displacement site at the foraging tree, nest site, Unfamiliar panorama test location (90° away from foraging site) and the Covered panorama (360° view-blocking at the foraging site). 360° images were taken with a Sony HD bloggie camera and converted into cylindrical images.

Control conditions were conducted at the same displacement sites on 75 foragers of the same nest that were not rewound (N = 25 in each condition). After their first inbound walk, individual ants were picked up before entering the nest and tested once in one condition (NPC, CPC or UPC). Testing procedures followed those for experimental (rewound) ants.

After testing, individual ants were caught and painted with black color to avoid the multiple testing of ants.

### Data analysis

We digitized all the hand-drawn recorded ant paths using GraphClick^TM^ into *x*-*y* co-ordinates and carried out further analysis in R Programming language. To determine the individual ant path straightness, the digitized co-ordinates from the release point to the end point at the nest were transferred into a R function (TrajFromcoords) that produces the trajectory of the ant.

Then to get the straightness index from the trajectory of the ant-path output, we measured the straight-line distance between the release point and end point of the path. The ratio of the straight-line distance from the displacement site to the end of the path to the total recorded path length (obtained by using another R function, TrajStraigtness) defined the straightness measure. The range of path straightness was between 0 to 1, with larger values indicating straighter paths (Islam, Deeti, Mahmuda, Kamhi, & Cheng, 2021b, Islam, Deeti, Murray, & Cheng, 2022). Scan rate (number of scans per meter of path traveled, defined by Wystrach et al., 2019), and initial heading directions of all ants in each trial were calculated.

On the displacement tests, both the manipulation tests and their corresponding control conditions, and on rewinding trials 1, 5, 9, the initial orientations were calculated and V, Rayleigh, and Watson tests performed by using the Oriana Version 4 (Kovach Computing Services^TM^) statistical program. Each bar in the circular plots is centered at the middle of the wedge that was crossed, with the length of the bar proportional to the frequency in the group. In order to check whether a distribution of headings was uniform, a Rayleigh test was performed. To evaluate whether initial orientations were significantly clustered around the nest direction at 0°, we examined whether 0° fits within the 95% confidence interval (CI) of orientations (Watson Tests) and performed V tests with alpha at *p* = 0.05. A V test yields a significant result when a heading distribution is significantly clustered around a given target direction. The mean directions of corresponding Treatment and Control distributions were compared with the parametric Watson-Williams multi-sample test.

### Data Transparency and Openness

Our data and R scripts for both Experiment 1 and Experiment 2 are made publicly available at (https://osf.io/na2uf/files/osfstorage). The predictions and methodologies for both experiments in this study were not pre-registered.

## Results

### Homeward run repetition after a single outbound run

We selected foragers that had completed a single outbound run on the evening before and let them run home from the release point before capturing them just before they entered their nest and displacing them back to the release point (Figure 1). Over repeated trials of rewinding, foragers were more variable in their trajectories, decreased their path straightness, increased the number of scans and also sometimes changed the initial heading direction after release (Figure 3A–3D, Figure 4, Supplementary Figure 1).

**Figure 3.**
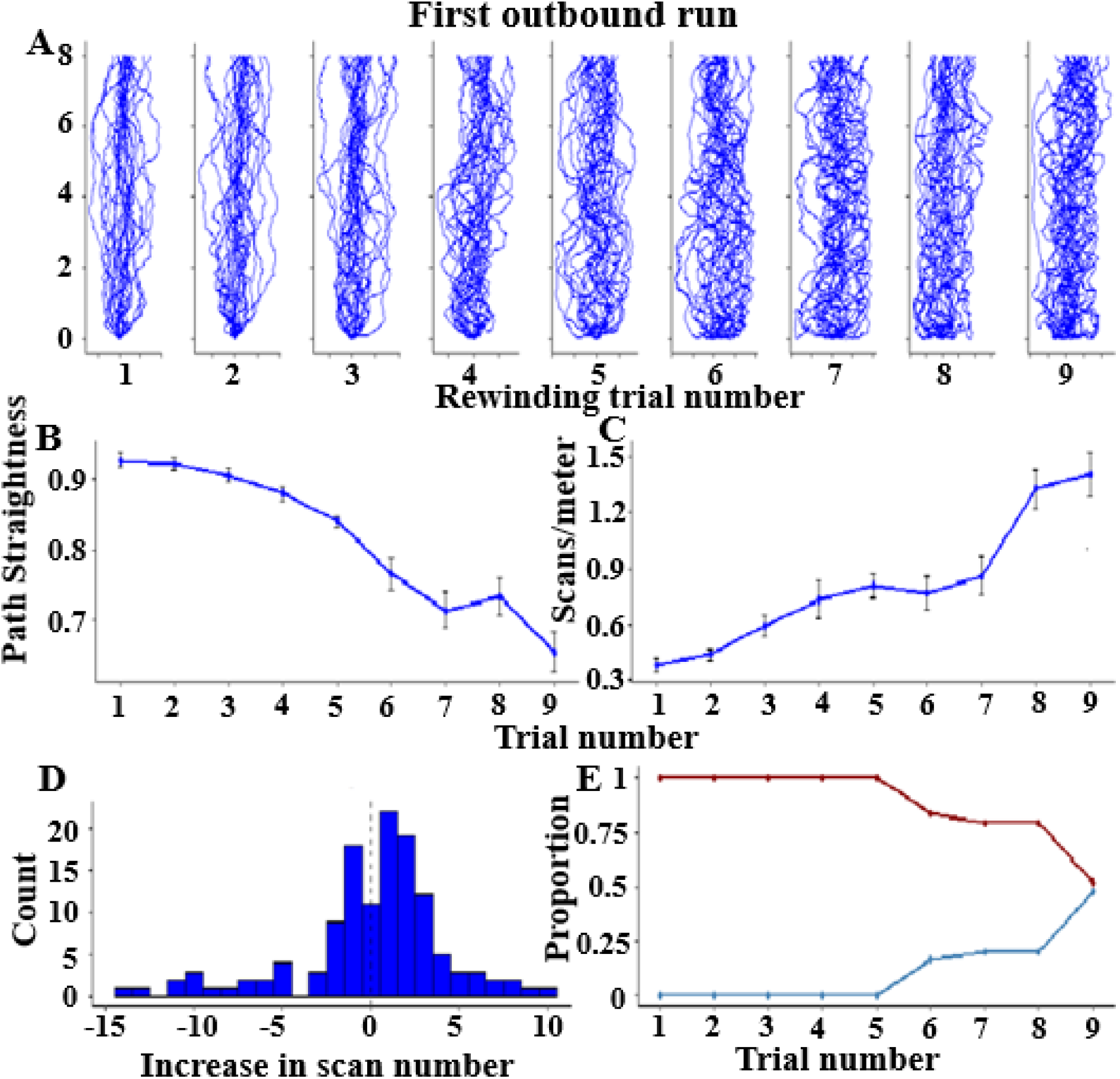
Complete rewinding paths and path parameters of foragers with repeated rewinding. (A) Paths of foragers in their 9 homeward reruns during the rewinding. (B) Mean path straightness and (C) mean scan rate per meter ± standard error of the mean for successive rewinding trials. (D) Changes in the scan rate from one run to the next. (E) Proportion of ants performing or not performing a U-turn on a homeward run.

**Figure 4.**
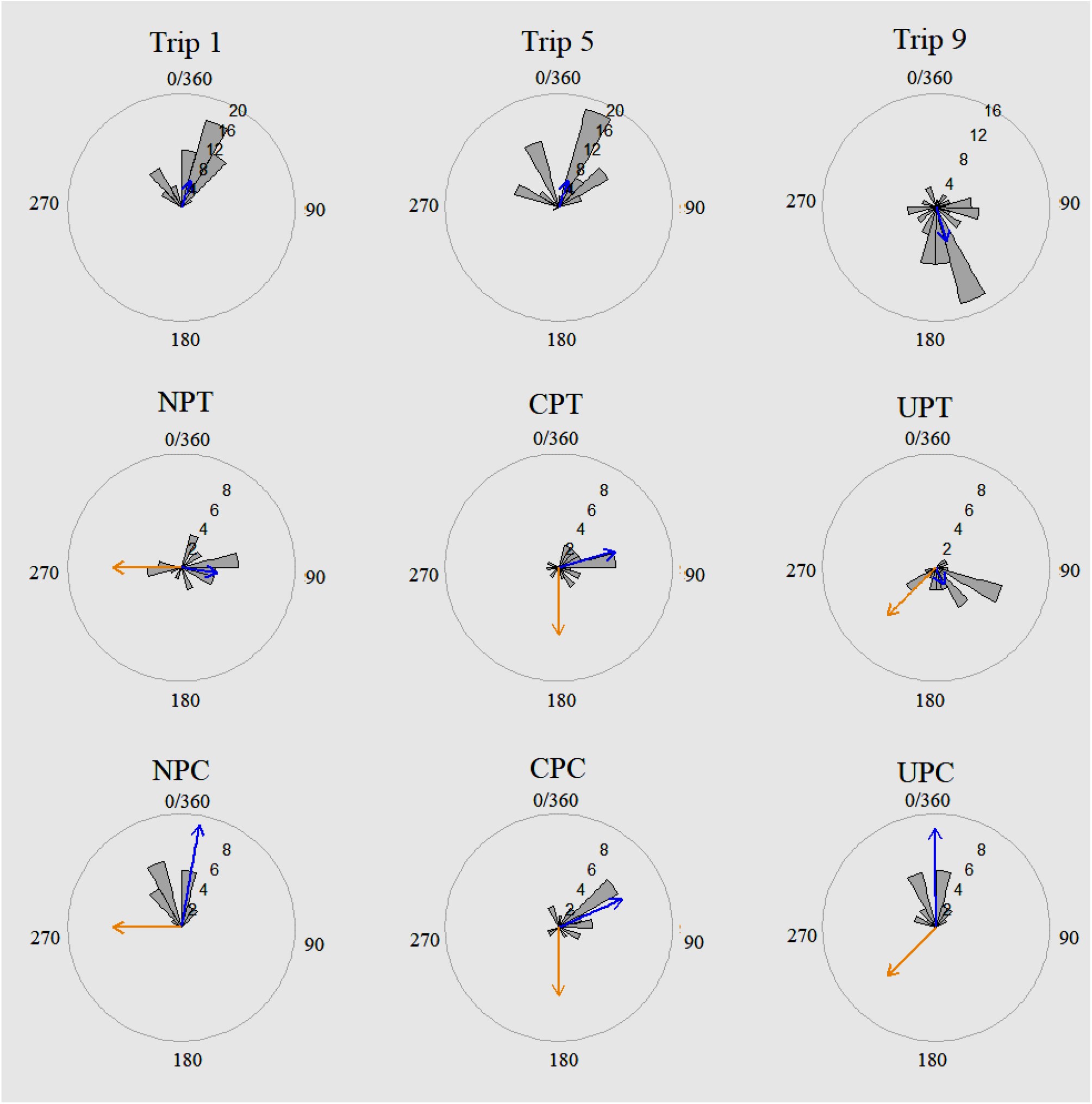
Circular histograms of initial headings of foragers during the homebound trips on rewinding trials 1, 5, and 9, and on tests with the Nest panorama (NPT), the Covered panorama (CPT), and the Unfamiliar panorama (UPT) and respective controls (NPC, CPC and UPC, in which ants were not rewound). In the histograms, the nest direction is set at 0°. The blue line segments denote the direction of the mean vector. The orange line segments denote rewinding vector direction (from nest to foraging tree).

Path straightness decreased significantly across trials (one-way ANOVA; *F*(8, 223) = 229.7, *p* < 0.001, linear trend: *F*(2, 222) = 229.7, *p* < 0.001, quadratic trend: *F*(2, 222) =118, *p* < 0.001), while the rate of scans increased significantly (one-way ANOVA, *F*(8, 127) = 119.8, *p* < 0.001, linear trend: *F*(2, 127) = 119.8, *p* < 0.001, quadratic trend: *F*(2, 127) = 63.56, *p* < 0.001). The histogram shows the change in scanning rate from the previous trial (Figure 3D; *p* <0.001 by a binomial test). The change in the number of scans from one run to the next run more often increased than decreased. The above-zero central tendency shows that scans increased with the number of re-runs. The linear mixed-model ANOVA shows a significant difference across the trials (*F*(8, 596) = 12.9, *p* = 0.005).

As a proxy for the point at which PI overrode the visually defined direction, we measured U-turn behavior, whereby ants started moving in the direction from the nest to the foraging tree (the direction of the accumulated PI vector, within ±22.5°; Figure 3E). U-turns first appeared during the 6th trial, and increased in proportion over subsequent trials. Once an ant U-turned, it always U-turned on all subsequent trials, which resulted in a survival curve of the proportion of the ants that did not perform a U-turn on each rewinding trial, with 50% of ants performing U-turns by walk 9 (Figure 3E). When ants took the U-turn, the path length in the opposite direction significantly increased with each rewinding trial (Supplementary Figure 2). The increased length as a proportion of the nest-to-foraging-tree distance increased from ∼6.52% in the 6^th^ rewinding trial to ∼7.06% in the 7^th^ rewinding trial to ∼8.20% in the 8^th^ rewinding trial up to ∼8.66% in the 9^th^ rewinding trial. As a proportion of the accumulated path-integration vector length (which increases in the nest-to-foraging-tree direction with each rewind), these proportions decreased slightly from rewinding trial 6 to rewinding trial 9: ∼1.49%, ∼1.34%, ∼1.34%, ∼1.24%, respectively. The data in Figure 3E show that even after 9 rewinding trips, some half of the foragers still did not take a U-turn. As described more fully below, however, few ants were oriented in the nest direction on rewinding trip 9. The ants that took a U-turn traveled initially only a short distance before turning back to head in the nest direction. Total path length, however, was affected by rewinding. The total path length traveled by the foragers from the displacement site to the nest (Figure 5) increased across rewinding trials (one-way ANOVA: *F*(8, 222) = 183.8, *p* < 0.001, linear trend: *F*(2, 222) = 183.8, *p* < 0.001, quadratic trend: *F*(2, 222)= 96.71, *p* < 0.001).

**Figure 5.**
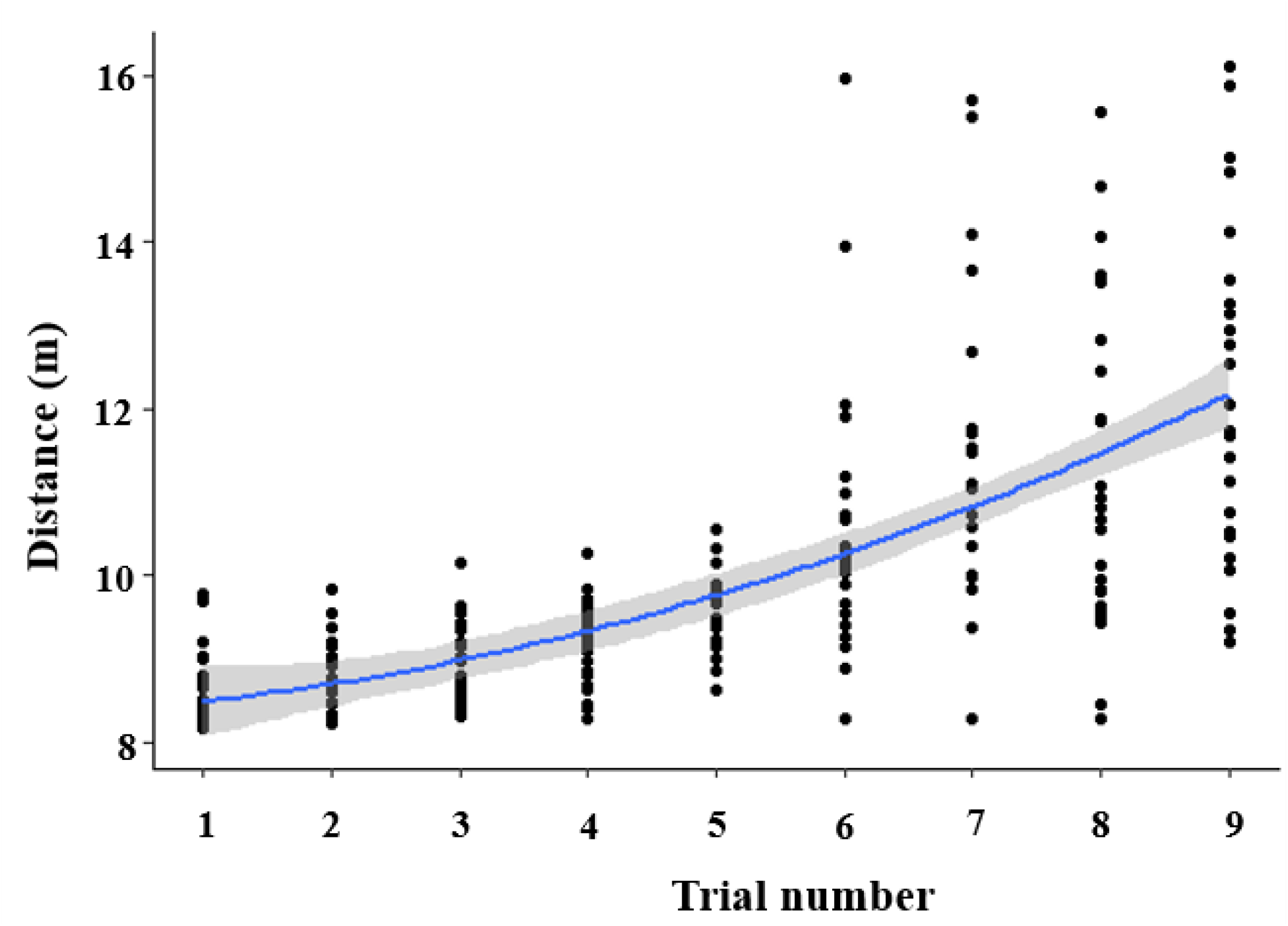
The total distance (path length) traveled by foragers during the rewinding trials. *Y*- values have been jittered on the plots. The total distance traveled by the foragers shows a quadratic trend. The blue line indicates a quadratic fit and the gray band shows the 95% confidence interval about the mean.

While the initial heading directions were nest-aligned in the earlier trials, by the last trial they were aligned in the opposite direction. The mean vector direction on trip 1 (*µ* = 10.51° and r = 0.91; V-test: *µ* = 10.76, p = 0.005) and trip 5 (*µ* = 12.54° and *r* = 0.77; V-test: *µ* = 9.13, p = 0.005; Figure 5) indicated that initial heading directions were oriented toward the nest direction (0°) while on the 9th trial (*µ* = 165.51° and *r* = 0.43), most of the initial orientations had shifted to the opposite nest-to-foraging-tree direction (180°) and few ants headed off toward the nest (Supplementary Figure 1). Ants were significantly oriented in the nest direction on trials 1 and 5 but not on trial 9 (V test, Table 1). On rewinding trial 9, the nest-to-foraging-tree direction condition (180°) showed significant V-test results (*µ* = 4.948, *p* = 0.005).

**Table 1:** Statistical results for initial heading directions in rewinding trips 1, 5, and 9, and on displacement tests after rewinding trial 9.

| **Test** | **RW** | **<u>Mean</u>**<br>**<u>vector</u>** | **<u>95% confidence</u>**<br>**<u>interval</u>** |  | **<u>Rayleigh test</u>** |  | **<u>V test detection 0<sup>0</sup></u>** |  |
| --- | --- | --- | --- | --- | --- | --- | --- | --- |
|  |  | **<u>μ</u>** | **<u>Minus</u>** | **<u>Plus</u>** | **<u>Z</u>** | **<u>P</u>** | **<u>Z</u>** | **<u>P</u>** |
| **Trip 1** | 0 | 10.51° | 4.38° | 16.64° | 59.92 | < 0.001 | 10.76 | < 0.001 |
| **Trip 5** | 4 | 12.54° | 3.14° | 21.6° | 43.82 | < 0.001 | 9.13 | < 0.001 |
| **Trip 9** | 8 | 165.51° | 145.51° | 185.5° | 14.21 | < 0.001 | -5.16 | 1 |
| **NP-T** | 9 | 98.81° | 50.12° | 147.5° | 2.52 | 0.07 | -0.34 | 0.63 |
| **CP-T** | 9 | 74.31° | 45.87° | 102.74° | 6.7 | < 0.001 | 0.99 | 0.16 |
| **UP-T** | 9 | 156.21° | 131.97° | 180.44° | 8.64 | < 0.001 | -3.8 | 1 |
| **NP-C** | 0 | 3.23° | 356.72° | 9.74° | 43.64 | < 0.001 | 9.32 | < 0.001 |
| **CP-C** | 0 | 65.98° | 40.73° | 91.24° | 7.79 | < 0.001 | 1.61 | 0.05 |
| **UP-C** | 0 | 359.58° | 347.75° | 11.41° | 18.91 | < 0.001 | 6.15 | < 0.001 |

Note. The nest direction for all conditions was at 0° with degrees. Increasing degrees
represent clockwise angular rotation. RW is the number of nest-bound trips completed. For
displacement tests, -T represents rewinding Treatment, while -C represents control ants that
had not been rewound. Locations are NP = Nest panorama, UP = Unfamiliar panorama, CP =
Covered panorama.

### Post-rewinding displacement tests

After the 9th trial, we subjected each forager to a single Post-rewinding treatment involving displacement to a specific location. As controls, we displaced un-rewound ants to each of these displacement locations. Rewound foragers displayed significantly different initial heading directions depending on which treatment they were subject to (Figure 4). At the Nest Panorama, displaced foragers’ headings formed a uniform distribution (Mean heading: *µ* = 98.81° and *r* = 0.318, Rayleigh: z = 2.52 *p* = 0.07). There appeared to be a tendency to walk along the foraging corridor, with some ants walking in the tree-to-nest direction (home direction) and others in the opposite direction (according to the accumulated PI vector direction), resulting in a potentially bimodal distribution. To test for a bimodal distribution we transformed the data by doubling each angle and found a non-uniform distribution (mean vector (*µ*) at = 223.79°; Rayleigh: *z* = 7.53, *p* = 0.001). This confirmed our suspicions since a transformed mean vector near 180° indicates bimodal peaks at ∼90° and ∼270° (V Test; *µ* = 2.114, *p* = 0.017). Control ants (which had not experienced rewinding) when tested at the Near Nest location were significantly nest-directed (*µ* = 3.23° and *r* = 0.91), which is significantly different from rewound ants (Watson-Williams test: *F*(1, 48) = 27.31 *p* = 0.0005). The Covered panorama treatment caused ants to head in an intermediate direction (*µ* = 74.31° *r* = 0.518; Rayleigh: *z* = 7.78, *p* = 0.001) between the foraging-tree-to-nest direction (0°) and its opposite, the nest-to-tree direction (180°). At the Unfamiliar displacement site, rewound ants were non-uniform in their heading, but not nest-directed (Rayligh: *z* = 8.64; *p* < 0.001, V-test: *z* = 3.8, *p* = 1) whereas unrewound ants were significantly directed toward the nest from here (*µ* = 359.58°, *r* = 0.87), which is significantly different from the initial directions of rewound ants (Watson-Williams test: *F*(1, 48) = 178.31 *p* = 0.0005). The Unfamiliar panorama location caused rewound foragers to head in a direction (*µ* = 156.21° and *r* =0.58) that was roughly 70° from the vector direction (∼228°). The rewound ants appear to head in the nest-to-foraging-tree direction at this Unfamiliar location (V-test results: *z* = –3.8, *p* < 0.001), although the 95% CI of the mean vector does not include this direction. Ants displaced in the Covered condition headed in a similar direction regardless of whether they were rewound or not (Treatment: *µ* = 74.31° and *r* = 0.51; Control: *µ* = 65.98° and *r* = 0.69; Figure 5, Table 1). The two distributions are not significantly different in mean heading (Watson-Williams test: *F*(1, 48) = 0.18 *p* = 0.66).

## Discussion

In summary, we found that successive rewinding interfered with foragers’ ability to orient to the true nest direction during their Initial headings, decreased their Path straightness, and resulted in increased scanning behaviors. This pattern mirrors the results of the rewinding experiment on the desert ant *Melophorus bagoti* (Wystrach et al., 2019). We then investigated the effect of rewinding on the ants’ navigational performance when views were altered. Near the nest, but directionally off of the usual route, ants did not head initially toward the nest, but in either the foraging-tree-to-nest compass direction or else the opposite compass direction, from the nest-to-foraging-tree, the latter of these orientations was consistent with the accumulated vector during rewinding. Control ants tested at this location without any rewinding homed readily to the true nest location, consistent earlier research (Freas et al., 2017). When we blocked the surrounding terrestrial visual cues (Covered panorama), ants chose an intermediate direction between the nest direction and its opposite. Control ants tested at this location without any rewinding performed similarly, so that rewinding did not change the ants’ behavior on this test. When displaced to an unfamiliar off-route panorama (Unfamiliar panorama), ants headed off initially roughly opposite to the nest compass direction; this direction was about 70° from the vector direction (nest-to-foraging-tree).

Control ants that had not been rewound headed in the nest compass direction from this location, showing that rewinding altered the ants’ behavior on this test. The rewinding trips completed by foragers showed that no foragers took a U-turn for the first 5 rewinding trips (6 trips including the initial trip), with all foragers following their homebound route to reach the nest. Beginning on trial 6, some ants began to exhibit U-turns, initially moving in the nest-to- foraging-tree direction (the vector direction) before turning and orienting towards their homebound route. The percentage of foragers which performed this behaviour increased with subsequent rewinds, yet even after 9 rewinding trials, ants that U-turned travelled only a short distance in the nest-to-foraging-tree direction, only traveling less than 2% of the vector length before abandoning this orientation.

With each additional rewinding, foragers travelling inbound to the nest continuously increased the length of the path integration vector, initially generating, then increasing the conflict between PI and visual information. Results on desert ants show that the conflict between the remembered landmarks and path integration contributes to path degradation (Wystrach et al., 2019), consistent with the current study. Both studies show that as the conflict between visual cues and path integration increases with rewinding runs, the path taken by the ants deteriorated, showing more meandering and scanning, and with more ants taking anti-nest headings (U-turns). The desert ants finished their experimental participation as soon as they U-turned (Wystrach et al., 2019), so that comparable data on the U-turn runs in *M. bagoti* are not available. Andel and Wehner’s (2004) *Cataglyphis* ants ran some 40% (maximal turning point, see their Figure 4a) of the vector distance in the U-turn direction after 7 rewinds, but in an environment (channel) with the usual experimental landmarks removed. Our U-turn results are consistent with the dominance of visual panoramic cues for this species (Freas et al., 2017). Why did our ants switch from vector-based navigation to view-based navigation? These navigational strategies in the ants’ toolkit work in parallel, with weights being adjusted ‘online’ (Hoinville & Wehner, 2018; Wehner et al., 2016). The weight accorded to vector navigation decreases as vector length decreases (Wystrach et al., 2015). But our ants traveled less than 2% of the vector length, making this factor improbable as a cause for switching strategy and turning back in the nest direction. *Myrmicia midas* resides in a visually rich and cluttered environment in which they rely heavily on terrestrial visual cues. Perhaps because of the visual complexity (see Figure 2), foragers took some time to process view-based information to determine the homeward direction. Work on bull ants has shown that ants are influenced by negative views or views to avoid as well as positive views or views to head toward (Le Möel & Wystrach, 2020, Murray et al., 2020). Such negative views might well have driven the course reversal found in the current study. We are suggesting that it might have taken a few seconds for the negative views to register and cause the ants to turn around.

The rewinding experiments on desert ants have shown that any change in the ants’ usual routine on their familiar routes typically leads to more meandering (decrease in path straightness) and pausing to scan along the path (Wystrach et al., 2019). Our interpretation is that these behaviors serve similar functions in the two species, to expose the navigator to different views of the environment, allowing them to discover new possibilities for reaching their nest. And like Wystrach et al. (2019), we also suggest that associative learning, likely supported by the mushroom bodies in insect brains, guides the ants’ travel by modulating the valence assigned to stimuli. It is likely that during rewinding, continued exposure to the homeward route, without the positive association of entering the nest, decreases (though does not eliminate) the positive memory valance associated with these inbound views. These lower positive memory valences then cause behaviour outputs such as more meandering and more interruptions of movement altogether to pause and scan, allowing ants, for example, to learn to detour around an obstacle or a trap (Wystrach et al., 2020).

Our experiments tested rewound ants when presented with different views, and these results provide further insights regarding the effects of rewinding. At the test site 1 m from the nest, but in a direction perpendicular to the foraging-tree-to-nest direction, the results showed that the ants were not oriented in the nest direction. Control ants that had not been rewound headed readily in the home direction, replicating earlier studies on this species (Freas et al., 2017; Freas & Cheng, 2019). Some rewound ants tested at the Nest panorama site headed in the vector direction. This outcome might be expected as about half did this on the immediate previous run (rewinding trial 9). But more ants headed initially in the foraging-tree-to-nest direction; the mean vector was in this direction, with the statistical analyses supporting a bimodal distribution of initial headings. This suggests that with rewinding, the ants had come to rely more on celestial cues rather than terrestrial views, at least for initial heading, which was all that was measured on the test. This reliance on celestial cues is surprising in that the ants did not use what must have been accurate and reliable terrestrial information to the ants, the view of their nest tree from 1 m away. Rewinding has changed behaviors even at locations that are familiar but not on the rewound route. Desert ants are known to recode the direction of travel from terrestrial to celestial cues when walking backwards (Schwarz, Mangan, Zeil, Webb, & Wystrach, 2017). Such an effect of rewinding and the topic of switching between terrestrial and celestial cues deserve further study (Reid et al., 2013).

When no useful views were available, rewound ants headed neither in the vector direction (something that they had done on the immediately preceding (9^th^) rewinding trial) nor in the foraging-tree-to-nest direction (Islam et al., 2022). The mean direction was toward the brighter portion of the sky at test time, so that the response might reflect positive phototactic behavior (Wehner, 1997). Control ants that had not been rewound behaved similarly on this test. We suggest that the change in context, from the view around the release point to the blocked view, had reduced the ants’ willingness to rely on the vector direction for initial orientation. A parallel phenomenon has been found in desert ants (*M. bagoti*) in that when the test context differs more from the context of their usual route, these ants travel a shorter distance in the vector direction before breaking off to engage in systematic search (Cheng, et al., 2012; Narendra 2007). When the view at the test site resembled that at the release point to some extent (on a wide-open field with only distant landmarks), the desert ants traveled about 80% of the vector distance, whereas when the view was very different (open field vs. cluttered landscape), the desert ants traveled only about 40% of the vector distance. Whatever the interpretation of the mean vector in this condition, the results show that a change in visual conditions has diminished the weight accorded to path integration, so that the ants no longer headed off in the vector direction as they did on the immediately preceding rewinding trial.

The results from the Unfamiliar panorama test might reflect some phototaxis as well. The mean direction of 156° lay somewhere between the vector direction (228°) and the direction of most light (∼125°). Rewinding played a role on this test as ants that had not been rewound headed readily in the home direction from this location. Once again, an unfamiliar view has weakened the propensity to use path integration, while rewinding likely weakened the propensity to use terrestrial information from this semi-familiar location.

Even when large conflicts are introduced to the path integration vector through rewinding, *M. midas* foragers rely predominantly on view-based homing. While after 9 rewinds, up to half of foragers initially headed in an anti-nest direction, they soon abandoned this PI orientation and regained a view-based nestward heading. Furthermore, the ease at which they maintain a nest-bound heading appears diminished by the procedure, as evidenced by less path straightness and more incidences of scanning behavior. Finally, rewinding appears to interfere with homing from locations away from the foraging corridor, with some evidence that compass information, rather than vector or visual information, determine heading choice, at least initially. In addition, the context-specific rewinding effects in these ants was not fully supported by the off-site test data.

### Experiment 2

The aim of Experiment 2 was to test context (view) specificity in the effects of rewinding by dividing the route home into two equal sections, then rewinding ants in only one of these sections. Each ant was rewound 4 times in its assigned section and then allowed to run the entire route home. We would expect that any effects of rewinding would be accentuated in the section in which the ants were rewound.

## Methods

In order to understand view specific effects during the ant navigation, the foraging corridor was divided into two sections representing half of the homeward route, Section-A (from the platform center to the midpoint home from there) and Section-B (from the midpoint to the nest). In rewinding in Section-A (n = 15), foragers were released at the center of the wooden platform and allowed to navigate 3.5 m of the straight-line distance from the displacement site toward the nest. The ants were collected in a dark graduated cylinder when they reached the edge of Section-A and returned to the starting point four times. On the fifth run, foragers were allowed to run the entire homeward path. We recorded the paths of the foragers, noting the number of scans and the locations of these scans during the rerunning walks. Similarly, a rewinding condition was performed in Section-B (n = 15). At the start of the 3.5m Section-B, ants were released and allowed to navigate through Section-B. Foragers were captured near the nest, returned to the same starting point, and allowed to re-run Section-B again three more times. On the 5th trial, ants were released on the wooden platform at a distance of 7 m from the nest and allowed to run the entire route home. We plotted the routes and noted the number of scans and the scans’ positions to examine view specificity in rewinding. View specificity or context specificity should be found in the form of interaction effects between the section being run on the 5th trial and the section in which an ant was rewound. We predicted that ants that were rewound in Section-A would exhibit higher deteriorations in their paths within Section-A than in Section-B. Conversely we would predict the opposite for ants rewound in Section-B: these foragers should exhibit higher uncertainty and higher path deterioration within Section-B than in Section-A.

### Data analysis

GraphClick^TM^ was used to digitize all recorded paths into *x*-*y* co-ordinates for further analysis. Scan rate (number of scans per meter of path traveled, defined by Wystrach et al., 2019) and path straightness (ratio of the straight-line distance from the displacement site to the end of the path to the total path length) of all ants in each trial were calculated. For analysis, we separated rewinding trials 1–4 from the 5th, test trial. A one-way repeated-measures ANOVA was used to determine if the means of path straightness and scan rate differed across rewinding trials 1–4, separately for rewinding in Section-A and in Section-B. Again, separating Section-A and Section-B rewinding trials 1 to 4, where ants crossed the 50 cm on the goniometers (initial orientations) were calculated and analysed as in Experiment 1. In order to probe context specificity in rewinding, we used a mixed model ANOVA on data from trial 5, when ants in both conditions (rewind in Section-A, rewind in Section-B) ran the entire route home. Scan rate and path straightness were the dependent variables examined.

Training section (rewind in Section-A, rewind in Section-B) was the between-subjects factor. Section run on the test (portion of path in Section-A, portion of path in Section-B) was the repeated-measures factor.

## Results

The initial headings of foragers from trip 1 to trip 4 and complete run in trip 5 were all significantly oriented toward the home direction in both Section-A and Section-B rewinding by the V test (Table 2, Figure 6, Supplementary Figure 2). Foragers’ path straightness showed a significant decrease over rewinding trials both in Section-A (One-way ANOVA, Section-A: *F*(3, 57) = 103.3, *p* = 0.001; linear trend, Section-A: *F*(3, 57)= 103.3, *p* = 0.001; quadratic trend, Section A: *F*(3, 57) = 52.21, *p* = 0.001), and in Section-B (One-way ANOVA, Section-B: *F*(3, 57) = 104.1, *p* = 0.001; linear trend, Section B: *F*(3, 57) = 104.1, *p* = 0.001; quadratic trend, Section B: *F*(3, 57) = 55.56, *p* = 0.001) ants. Scan rates increased across rewinding trials. Foragers scanned more frequently as the homebound trips increased (Section-A: *F*(3, 70) = 32.1, *p* = 0.001; Section-B: *F*(3, 70) = 20.91, *p* = 0.001).

**Figure 6.**
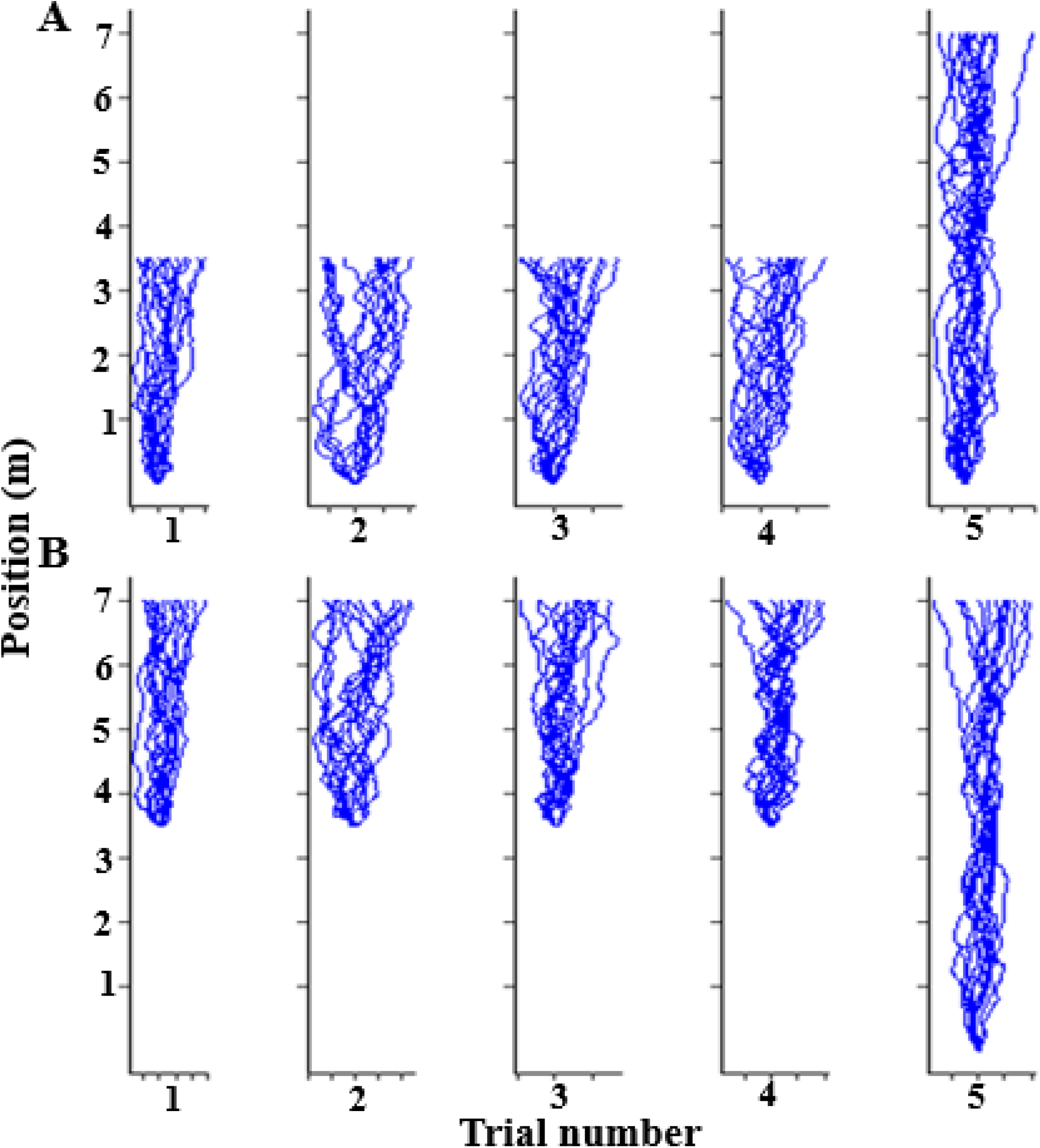
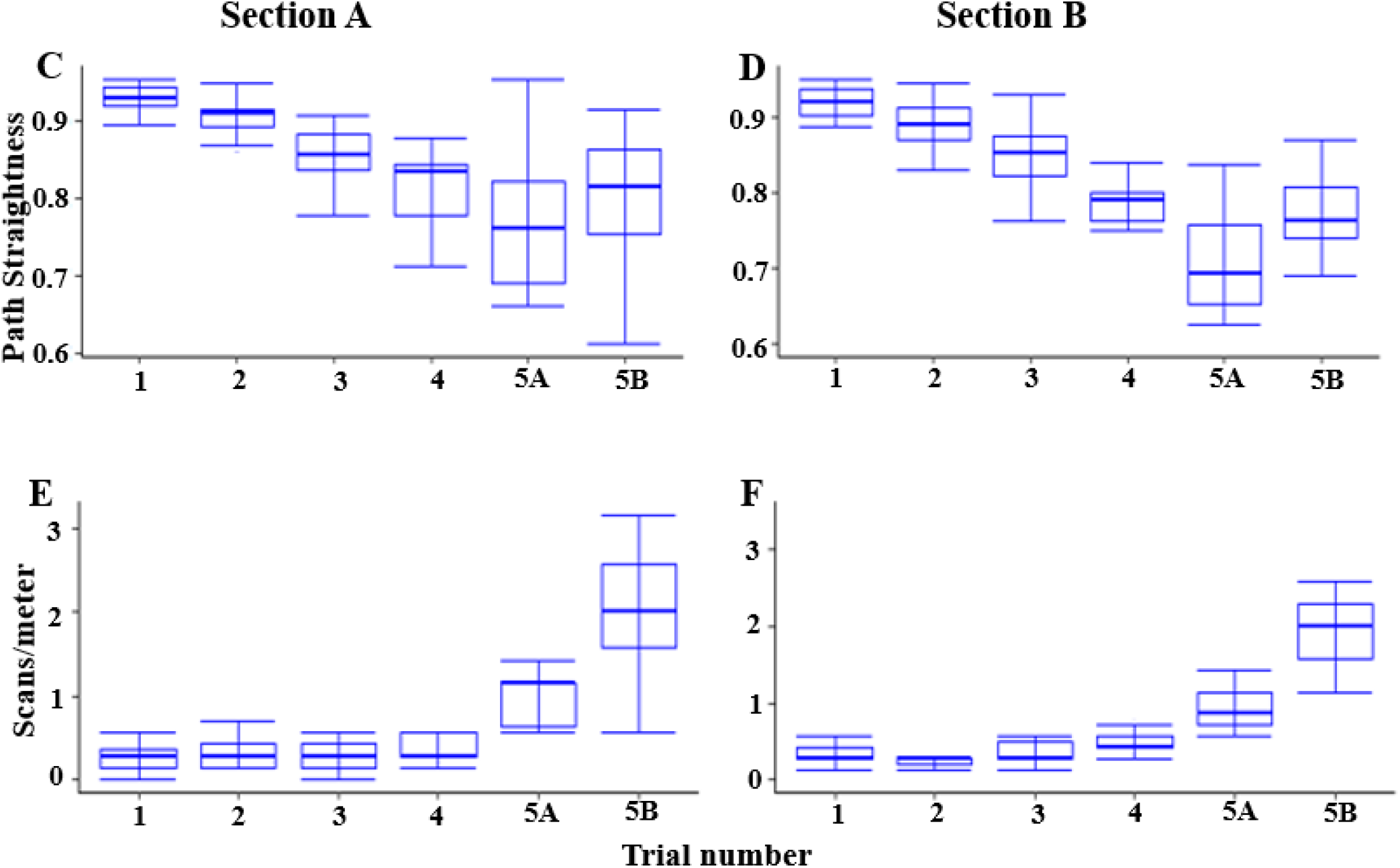
Rewinding paths in Section-A (A), Section-B (B), path straightness in Section-A rewinding runs (C) and in Section-B rewinding runs (D), and scans per meter in Section-A rewinding runs (E) and in Section-B rewinding runs (F). Ants were rewound in the 1st half or 2nd half of their route home for 4 times, and on the 5th trial ants walked the complete route home. The box plots show the median (middle line in the box), the lower and upper quartiles of the box. The whiskers go from the minimum to the maximum.

**Table 2:**
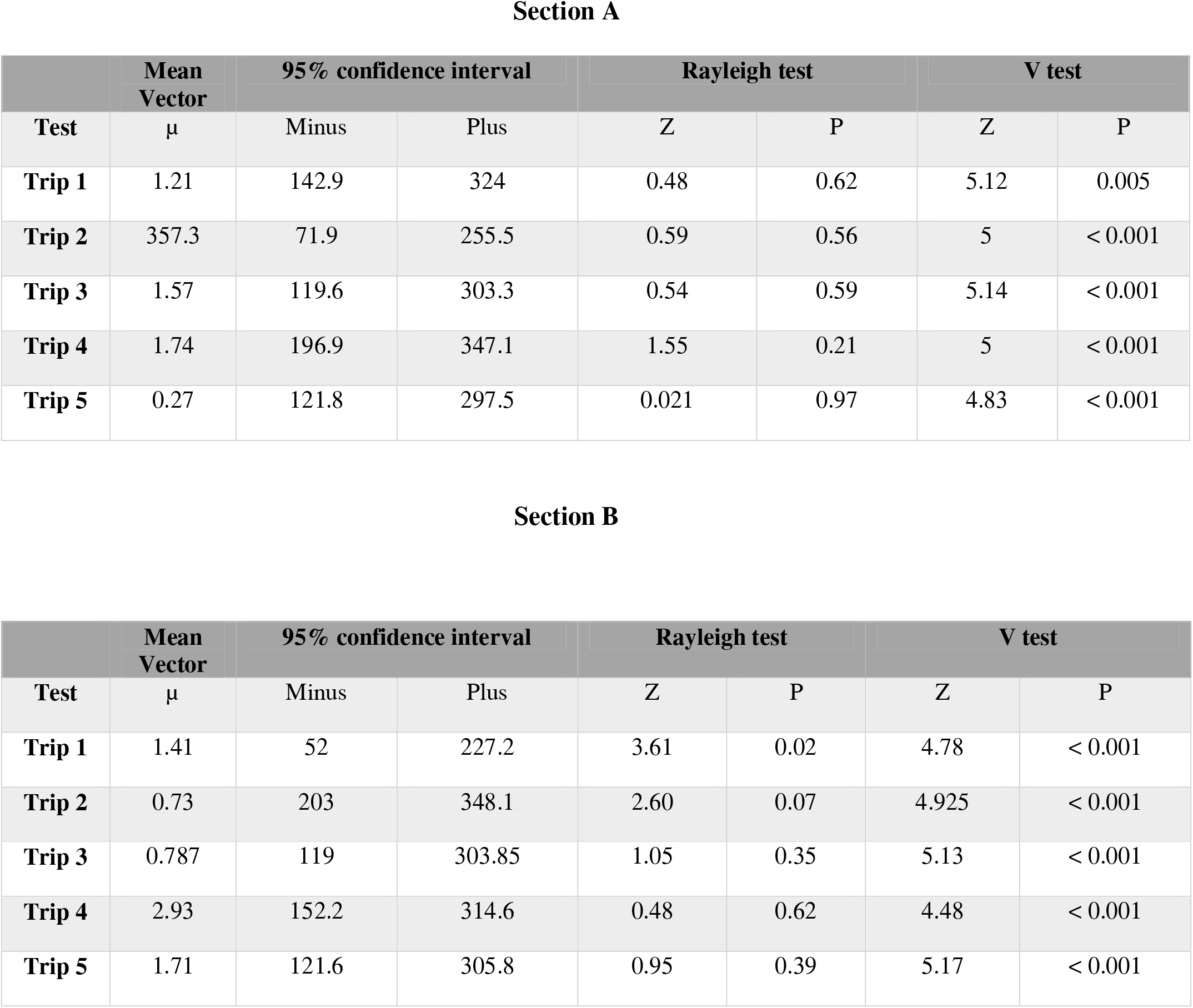
Statistical results for initial heading directions in rewinding trips of Section-A and Section- B 1 to 5 trials.

When we examined the 5^th^ and final full trip, we found significant effects of Treatment and Section on path straightness, but only Section had a significant effect on the number of scans. The ants rewound in Section-A had significantly straighter paths than those rewound in Section-B, (treatment main effect: *F*(1, 42) = 6.52, *p* = 0.014). In addition, paths were straighter closer to the nest (in Section-B) (section main effect: *F*(1, 42) = 7.25, *p* = 0.01), but no significant interaction between Treatment and Section (*F*(1, 42) = 0.42, *p* = 0.52) was found. By contrast we found no significant effect of Treatment on scanning (*F*(2, 56) = 0.04, *p* = 0.83), but, as with path straightness, ants scanned more in Section-B than in Section-A (*F*(2, 56) = 42.8, *p* < 0.001). There was also no interaction between Treatment and Section on scanning (*F*(2, 56) = 0.41, *p* = 0.52).

## Discussion

In Experiment 2, foragers were rewound only in the first half or last half of the foraging corridor to test for view specificity in the effects of rewinding. Just as in Experiment 1, forager’s path straightness was significantly reduced as rewinding trials increased. Scan rate increased over the trials, and more scans were found in Section-B. When the fifth run of the entire homeward path was analysed, however, we found no interaction between treatment and Section in both path straightness or in scans. Ants did not exhibit higher uncertainty or more disrupted in their homing behaviour within the section in which they were rewound.

In contrast, desert ants showed more path disruption in the section in which they were rewound (Wystrach et al., 2019). Unlike desert ants, view specificity in the effects of rewinding was not found in *M. midas*. Instead, foragers’ paths deteriorated throughout the route in current experiment. Why? A few possibilities can be considered but at this point, the answer remains uncertain until further investigations are made. The two species have different phylogenetic histories and the difference may be a matter of their respective phylogenetic signatures. That is, they belong to clades in which this trait (view specificity of rewinding) is present or not. This hypothesis can only be tested by examining more species on this trait. The two species have different lifestyle characteristics. *M. midas* workers live much longer than *M. bagoti* foragers do, and *M. midas* is night-active, whereas *M. bagoti* is day-active. Another possible confound is that this species of ants, from personal observations, scan more in general when they near their nest. This propensity might have dominated over any context-specificity effect. In the study on *M. bagoti* (Wystrach et al., 2019), section B ended 3 m from the ants’ nest and any increase in scans near the nest would not have been recorded in these ants. An additional alternative interpretation of these findings, that the results of Experiment 1 suggest, is that 9 rewinds produced more substantial effects than 4 and that contextual specificity might require a more extensive rewind manipulations to produce detectable differences along the route. To our mind, the most likely explanation stems from differences in the testing set ups in our study and Wystrach et al. (2019).

Wystrach et al. added experimentally provided landmarks and even obstacles such as baffles along the route home. The current study simply used the natural landscape, which while visually complex, does not contain many local landmarks that make route sections highly distinct. As a result, the two sections selected for comparison were far more distinct in Wystrach et al.’s study than in ours. These visual route differences lead us to believe that our tested ants were probably more likely to generalize their rewinding experience to the entire route. Of course, it would take further research to confirm or rule out such a hypothesis.

## Summary

Rewinding manipulations test how ants resolve diametrically opposing conflicts between cue sets, the path integration vector length and the visual scene around them. In these nocturnal bull ants, the visual scene won the conflict. Over successive rewinding trials, the bull ants kept on using the visual scene for homing, another demonstration of the importance of view-based homing in this species. Even if ants headed off initially in the opposite direction, the travel was cut short, and ants turned back to head in the nest direction. Repeated rewinding, however, led to path deteriorations, with more meandering and scanning, as it does in desert ants. Surprisingly, the effects of rewinding were not view specific. The bull ants did not deteriorate more in the section of space in which they were rewound.

## Acknowledgements

We thank Macquarie University for giving us support and access to the field site on campus. We acknowledge the traditional custodians of the land on which the field site sits, the Wallumattagal clan of the Dharug Nation. We acknowledge and pay our respects to Elders past and present.

## Author contributions

Experiments conceived and designed by SD and KC. SD and MI carried out experimentation and collected data. SD and TM analysed the data. SD, CF, TM and KC drafted and revised the manuscript.

## Ethics standards

Australia has no ethical regulations regarding work with insects. The study was non-invasive and no long-term aversive effects were found on the nests or on the individuals studied.

## Data availability

Excel files and R scripts are available at Open Science framework: https://osf.io/na2uf/files/osfstorage

